# A High-Throughput Extraction and Analysis Method for Steroidal Glycoalkaloids in Tomato

**DOI:** 10.1101/2019.12.23.878223

**Authors:** Michael P. Dzakovich, Jordan L. Hartman, Jessica L. Cooperstone

## Abstract

Tomato steroidal glycoalkaloids (tSGAs) are a class of cholesterol-derived metabolites uniquely produced by the tomato clade. These compounds provide protection against biotic stress due to their fungicidal and insecticidal properties. Although commonly reported as being anti-nutritional, both *in vitro* as well as pre-clinical animal studies have indicated that some tSGAs may have a beneficial impact on human health. However, the paucity of quantitative extraction and analysis methods presents a major obstacle for determining the biological and nutritional functions of tSGAs. To address this problem, we developed and validated the first comprehensive extraction and UHPLC-MS/MS quantification method for tSGAs. Our extraction method allows for up to 16 samples to be extracted simultaneously in 20 minutes with 93.0 ± 6.8% and 100.8 ± 13.1% recovery rates for tomatidine and alpha-tomatine, respectively. Our ultra-high-performance liquid chromatography tandem mass spectrometry (UHPLC-MS/MS) method was able to chromatographically separate analytes derived from 16 tSGAs representing 9 different tSGA masses, as well as two internal standards, in 13 minutes. Tomato steroidal glycoalkaloids that did not have available standards were annotated using high resolution mass spectrometry as well as product ion scans that provided fragmentation data. Lastly, we utilized our method to survey a variety of commonly consumed tomato-based products. Total tSGA concentrations ranged from 0.7 to 3.4 mg/serving and represent some of the first reported tSGA concentrations in tomato-based products. Our validation studies indicate that our method is sensitive, robust, and able to be used for a variety of applications where concentrations of biologically relevant tSGAs need to be quantified.

## 1 Introduction

Solanaceous plants produce a spectrum of cholesterol derived compounds called steroidal glycoalkaloids. While each solanaceous clade produces its own unique assortment of steroidal glycoalkaloids, these metabolites share commonality in their role as phytoanticipins and anti-herbivory agents (Etalo et al., 2015; Fontaine et al., 1948; Irving et al., 1945; Ökmen et al., 2013). Tomato (*Solanum lycopersicum* and close relatives) is no exception, and over 100 tomato steroidal glycoalkaloids (tSGAs, Fig. 1) have been suggested (Iijima et al., 2013, 2008). Although these compounds are typically reported as anti-nutritional (Ballester et al., 2016; Cárdenas et al., 2016, 2015; Itkin et al., 2013), other studies suggest a health-promoting role. In fact, emerging evidence suggests that some tSGAs may play a role in positive health outcomes associated with tomato consumption (Cayen, 1971; Choi et al., 2012; Cooperstone et al., 2017; Lee et al., 2004). While these compounds continue to be evaluated both *in planta* and *in vivo*, there is a lack of quantitative and validated methods to extract and measure tSGAs from tomatoes; a critical need for additional research in this area.

**Fig. 1.**
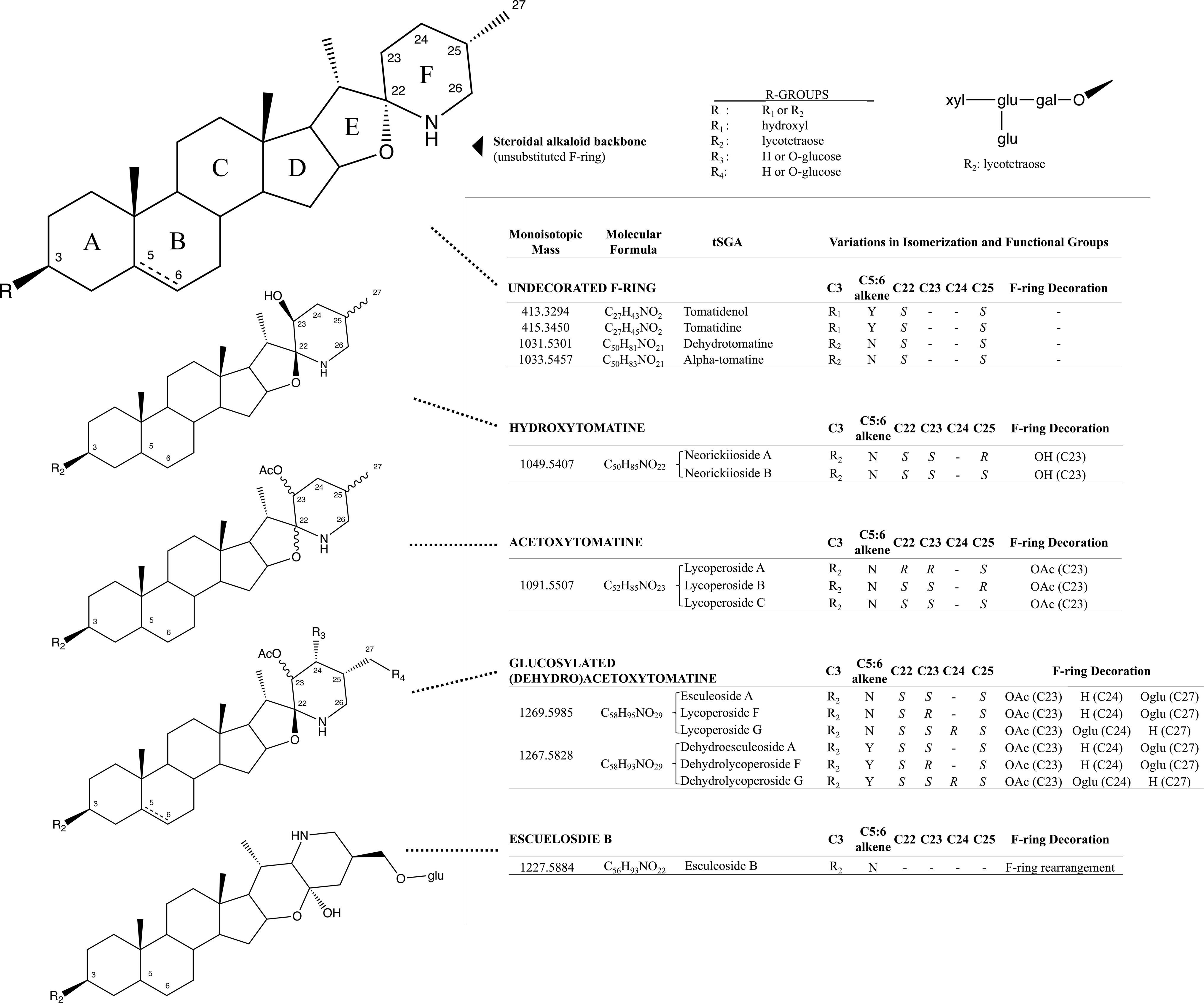
Structural and isomeric variation in selected tomato steroidal alkaloids. Steroidal glycoalkaloids found in tomato (tSGAs) are spirosolane-type saponins with variations in a singular double-bond (C5:6), F-ring decorations (C22-C27), F-ring rearrangement (resulting in a change in stereochemistry at C22), and C3 glycosylation (typically a four-sugar tetrasaccharide, lycotetraose). The undecorated SA backbone is shown first with relevant carbons numbered and ring names (A-F). Steroidal alkaloids (SAs) were grouped based on structural similarity with bonds of varying stereochemistry denoted by wavy bonds and varying C5:6 saturation status denoted by a dashed bond. Structural variation, along with the monoisotopic mass, molecular formula, and common name are displayed alongside structures for each group. R-groups were used to denote status of C3 glycosylation in all groups (R_1_ and R_2_) and possible positions of glucosylation on glucosylated (dehydro)acetoxytomatine (R_3_, R_4_). All possible isomers and derivatives are not shown, just those quantitated in this method.

Tomato steroidal glycoalkaloids are typically extracted by grinding individual samples using a mortar and pestle, or blender and then solubilizing analytes with polar solvent systems, typically methanol. This approach is time consuming because each sample is handled individually. Additionally, this technique has been used for relative profiling, and has not been evaluated for its ability to extract tSGAs quantitatively. Tomato steroidal glycoalkaloids such as alpha-tomatine, have previously been quantified using gas and liquid chromatography (Kozukue and Friedman, 2003; Lawson et al., 1992; Rick et al., 1994), as well as a number of bioassays including cellular agglutination (Schlösser and Gottlieb, 1966) and radioligand assays using radioactive cholesterol (Eltayeb and Roddick, 1984). These methods are unreliable, suffer from poor sensitivity, have poor selectivity for different alkaloids, and are time consuming. Recent advances in analytical chemistry have enabled researchers to discover other tSGA species in tomato fruits using high resolution mass spectrometry (Iijima et al., 2013, 2008; Zhu et al., 2018), however these methods are qualitative. A small number of quantitative methods using mass spectrometry have been developed, but only for individual or few of tSGAs (Baldina et al., 2016; Caprioli et al., 2014). Thus, there is a need to develop validated extraction and quantification methods in order to continue to study the role these compounds have in both plant and human health.

To address the lack of suitable approaches to extract and quantify tSGAs, we developed and validated a high-throughput extraction and ultra-high-performance liquid chromatography tandem mass spectrometry (UHPLC-MS/MS) method suitable for tomato and tomato-based products. Our extraction method is able to process 16 samples in parallel in 20 minutes (1.25 min/sample) and our UHPLC-MS/MS method can chromatographically separate, detect, and quantify 16 tSGAs (using two external and two internal standards) representing 9 different tSGA masses (Fig. 2) in 13 minutes per sample. This is the first comprehensive targeted method to quantify a broad panel of tSGAs. Here, we present the experiments used to develop and validate our method as well as an application providing baseline information of tSGA concentrations in commonly consumed tomato products.

**Fig. 2.**
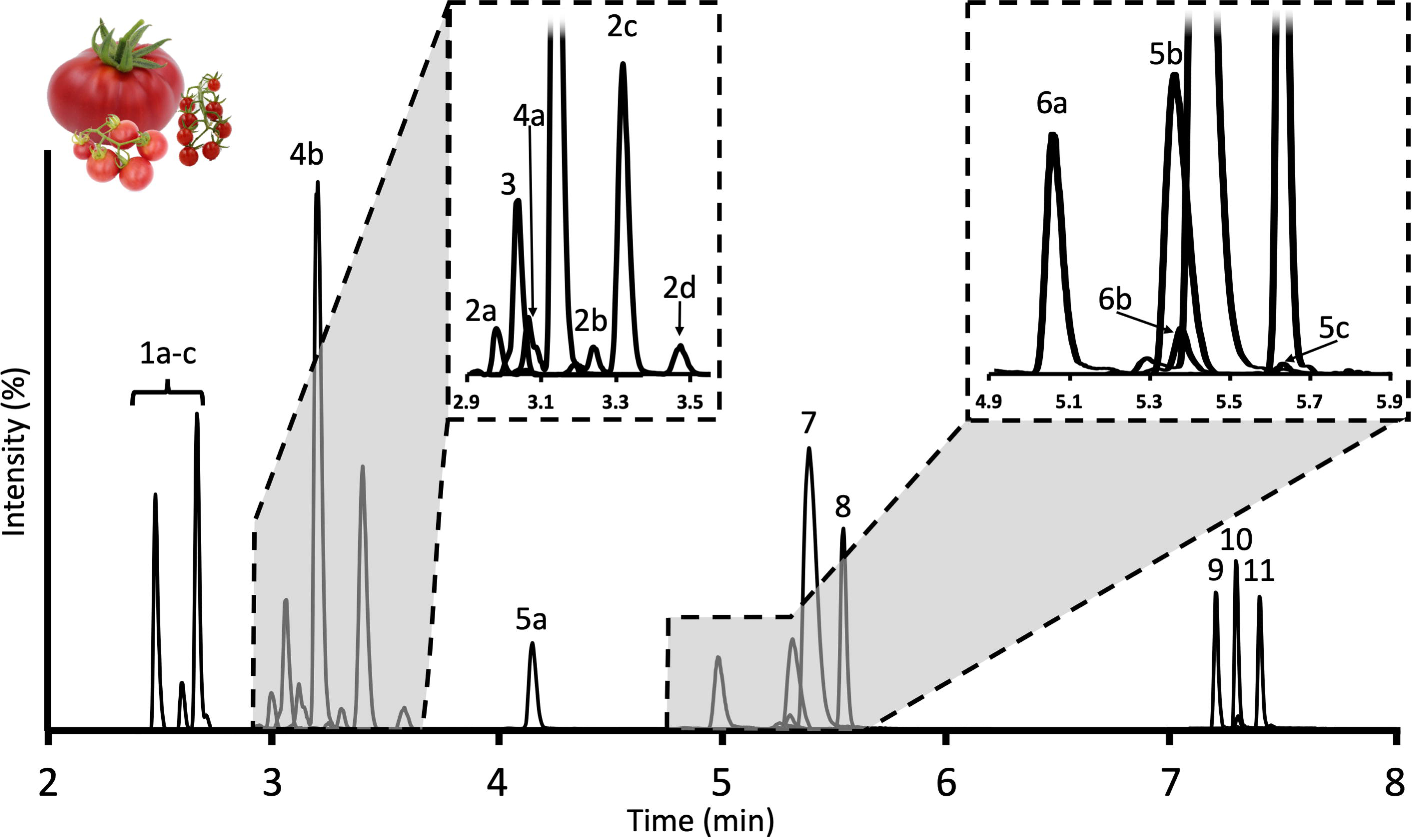
Chromatogram of tSGAs found in red ripe tomatoes measured by our UHPLC-MS/MS method. Peaks are identified as follows: **1a-c**: Esculeoside B1-3; **2a-d**: Hydroxytomatine; **3**: Dehydrolycoperoside F, G, or Dehydroesculeoside A; **4a**,**b**: Lycoperoside F, G, or Esculeoside A; **5a-c**: Acetoxytomatine; **6a**,**b**: Dehydrotomatine; **7**: Alpha-tomatine; **8**: Alpha-solanine; **9**: Solanidine; **10**: Tomatidine; **11**: Tomatidenol

## 2 Materials and Methods

### Reagents and standards

Acetonitrile (LC-MS grade), formic acid (LC-MS grade), isopropanol (LC-MS grade), methanol (HPLC grade), and water (LC-MS grade) were purchased from Fisher Scientific (Pittsburgh, PA). Alpha-tomatine (≥90% purity) and solanidine (≥99% purity) were purchased from Extrasynthese (Genay, France). Alpha-solanine (≥95% purity) and tomatidine (≥95% purity) were purchased from Sigma Aldrich (St. Louis, MO). Stock solutions were prepared by weighing each analyte into glass vials and dissolving into methanol prior to storage at −80 °C. Standard curves were prepared by mixing 15 nmol of alpha-tomatine and 1 nmol of tomatidine in methanol. The solution was evaporated to dryness under a stream of ultra-high purity (5.0 grade) nitrogen gas. The dried residue was then resuspended in 900 µL of methanol, briefly sonicated (∼ 5 s), and then diluted with an additional 900 µL of water. An 8-point dilution series was then prepared, and analyte concentrations ranged from 3.81 pmol/mL to 8.34 nmol/mL (11.14 femtomoles to 25 picomoles injected).

To utilize alpha-solanine and solanidine as internal standards (IS), 1.25 nmol and 22.68 pmol of alpha solanine and solanidine, respectively, were spiked into each vial of the alpha-tomatine/tomatidine external standard dilution series described above. The spike intensity of alpha-solanine and solanidine was determined by calculating the amount needed to achieve target peak areas of tSGAs typically seen in tomato samples.

### Sample material

For UHPLC-MS/MS and UHPLC-Quadrupole Time-of-Flight Mass Spectrometry (UHPLC-QTOF-MS) method development experiments, 36 unique accessions of tomato including *Solanum lycopersicum, Solanum lycopersicum* var. *cerasiforme*, and *Solanum pimpinellifolium* were combined and pureed to create a tomato reference material expected to span the diversity of tSGAs reported in nature. For spike-in recovery experiments, red-ripe processing-type tomatoes (OH8245; courtesy of David M. Francis) were diced, mixed together by hand, and stored at −20 °C until analysis. Items used for the tomato product survey were purchased from supermarkets in Columbus, OH in July 2019. Three unique brands of tomato paste, tomato juice, diced tomatoes, whole peeled tomatoes, ketchup, pasta sauce, and tomato soup were analyzed for tSGAs. Additionally, four heirloom, two fresh-market, one processing, and one cherry variety of unprocessed tomatoes were also analyzed.

### Extraction of tSGAs

Five grams of diced OH8245 tomato (± 0.05 g) were weighed in 50 mL falcon tubes. Two 3/8” x 7/8” angled ceramic cutting stones (W.W. Grainger: Lake Forest, IL; Item no.: 5UJX2) were placed on top of the tomato sample and 100 µL of internal standard was added, followed by 15 mL of methanol. Samples were then extracted for 5 minutes at 1400 RPM using a Geno/Grinder 2010 (SPEX Sample Prep: Metuchen, NJ). Sample tubes were immediately centrifuged for 5 minutes at 3000 x *g* and 4 °C. Two mL aliquots of supernatant from each sample were then transferred to glass vials and diluted with 1 mL of water. Samples were then filtered into LC vials using a 0.22 µm nylon filter (CELLTREAT Scientific Products: Pepperell, MA).

Tomato products sourced from grocery stores were extracted as described above except fresh fruits of each type were blended in a coffee grinder prior to extraction. To account for differences in water content among the tomato products, 500 µL aliquots from each sample were dried down under nitrogen gas, re-dissolved in 1.5 mL of 50% methanol, and filtered using a 0.22 µm filter prior to analysis.

### UHPLC-MS/MS Quantification of tSGAs

Tomato steroidal glycoalkaloids were chromatographically separated on a Waters (Milford, MA) Acquity UHPLC H-Class System using a Waters C18 Acquity bridged ethylene hybrid (BEH) 2.1 × 100 mm, 1.7 µm particle size column maintained at 40 °C. The autosampler compartment was maintained at 20 °C. A gradient method with Solvent A (water + 0.1% (*v*/*v*) formic acid) and Solvent B (Acetonitrile + 0.1% (*v*/*v*) formic acid) at a flow rate of 0.4 mL/min was utilized as follows: 95% A for 0.25 minutes, 95% A to 80% A for 1.0 minute, 80% A to 75% A for 2.5 minutes, 75% A held for 0.5 minutes, 75% A to 68% A for 1.7 minutes, 38% A to 15% A for 1.7 minutes, 0% A held for 3.0 minutes, and back to 95% A for 2.35 minutes to re-equilibrate the column. Each run lasted 13 minutes and the sample needle was washed for 10 seconds with 1:1 methanol:isopropanol before and after each injection to minimize carryover. Column eluent was directed into a Waters TQ Detector tandem mass spectrometer and source parameters and transitions can be found in Table 1. Dwell times were optimized for each analyte to allow for 12-15 points across each peak. Quantification was carried out using 6-8 point external calibration curves, depending on the extent of linearity for a given analyte. Relative quantification was used for tSGAs (quantified using alpha-tomatine) and their aglycones (quantified using tomatidine) that did not have commercially available standards. Additionally, signals were normalized to alpha-solanine and solanidine for glycosylated and aglycone analytes to correct for instrument variability.

**Table 1.** LC-MS/MS MRM parameters of steroidal glycoalkaloids quantified by our method.

| Analyte | Retention Time<br>(min) | Parent Mass<br>[M+H]<sub>a</sub> | Product Ions | Cone<br>Voltage (V) | Collision<br>Energy (eV) |
| --- | --- | --- | --- | --- | --- |
| Esculeoside B | 2.55, 2.67, 2.74 | 1228.6 | 254.9*, 1048.8 | 75 | 65, 40 |
| Hydroxytomatine | 3.02, 3.28, 3.37,<br>3.57 | 1050.6 | 254.9, 1032.7 | 55 | 55, 30 |
| Dehydrolycoperoside F, G, or<br>Dehydroesculeoside A | 3.08 | 1268.6 | 252.9*, 1208.9 | 80 | 65, 35 |
| Lycoperoside F, G, or Esculeoside A | 3.11, 3.26 | 1270.6 | 1048.8, 1210.9 | 70 | 60, 30 |
| Acetoxytomatine<sub>b</sub> (I) | 4.28 | 1092.6 | 84.7, 1032.7 | 40 | 65, 35 |
| Dehydrotomatine<sub>c</sub> | 5.09, 5.49 | 1032.5 | 84.7, 252.9* | 70 | 80, 50 |
| Acetoxytomatine (II) | 5.42, 5.66 | 1092.6 | 144.7, 162.8 | 40 | 50, 45 |
| Alpha-tomatine<sub>c</sub> | 5.45 | 1034.6 | 84.7*, 160.8 | 70 | 85, 60 |
| Alpha-solanine<sub>c,d</sub> | 5.64 | 869.1 | 97.8*, 399.1 | 70 | 85, 65 |
| Solanidine<sub>c,d</sub> | 7.22 | 398.7 | 80.7, 97.8* | 70 | 55, 35 |
| Tomatidine<sub>c</sub> | 7.30 | 416.4 | 160.8, 254.9 | 50 | 30, 30 |
| Tomatidenol<sub>c</sub> | 7.36 | 414.3 | 125.8, 270.7 | 40 | 30, 20 |

<sub>a</sub>Analytes were quantified using the following settings: Mass span: 0.3 Da, Capillary voltage: 0.5
kV, extractor voltage: 5 V, RF Lens voltage: 0.5 V, source temperature: 150°C, desolvation
temperature: 500°C, desolvation flow rate: 1000 L/hr, cone gas flow rate: 50 L/hr.
<sub>b</sub>Commonly referred to as lycopersides A, B, or C
<sub>c</sub>Indicates that analyte was confirmed by authentic standard
<sub>d</sub>Indicates analyte used as an internal standard.
\*Indicates quantifying ion; other ions used for qualifying purposes. Compounds with no indicated quantifying ion were quantified using the sum of both MRM transitions.

### UHPLC-QTOF/MS Confirmation of tSGA Identities

We verified the identities of our tSGA analytes using an Agilent 1290 Infinity II UHPLC coupled with an Agilent 6545 QTOF-MS. Identical column and chromatographic separation conditions were used as described above for our MS/MS method. The QTOF-MS used an electrospray ionization source operated in positive mode and data were collected from 50-1700 *m/z* for both full-scan and MS/MS experiments. Gas temperature was set to 350 °C, drying gas flow was 10 L/min, nebulizer gas flow was 10 L/min, nebulizer was 35 psig, and sheath gas flow and temperature was 11 L/min and 375 °C, respectively. For MS/MS experiments on the QTOF-MS, identical parameters were used except for the selection of tSGA masses of interest and a two-minute retention time window around each analyte to maximize duty cycle of the instrument. Collision energy for all tSGAs was set to 70 eV and all aglycones were fragmented with 45 eV.

### Limit of Detection (LOD) and Limit of Quantification (LOQ)

Limit of detection and LOQ were calculated using six replicates of the lowest concentration standard curve calibrant sample (3.81 and 0.254 femtomoles on column for alpha-tomatine and tomatidine, respectively) and determining their signal to noise ratios. Moles on column at 3/1 and 10/1 signal to noise were then determined for alpha-tomatine and tomatidine to calculate LOD and LOQ.

### Spike Recovery Experiments

Ten, 5 g (± 0.01 g) replicates of diced OH8245 processing tomatoes were weighed into 50 mL falcon tubes. Five samples were extracted as outlined previously with the addition of a 100 µL methanolic solution containing 1.67 nmol of alpha-tomatine, 1.25 nmol of alpha solanine, 12.4 pmol of tomatidine, and 22.68 pmol of solanidine (spiked tomato) while another five samples were extracted without IS solution (non-spiked tomato; 100 µL of methanol used in its place). The IS was allowed to integrate into the sample matrix for 30 min. Another set of five samples were prepared by substituting tomato for 5 mL of water for tomato and extracted with the addition of 100 µL of the methanolic IS solution mentioned previously (spiked mock sample). Percent recovery was estimated using the following equation:

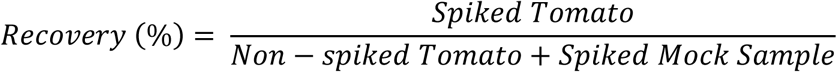

### Intra/Interday Variability Experiments

Eight OH8245 tomato fruits were blended together and 5 g aliquots (± 0.05 g) were distributed among 18, 50 mL tubes, and frozen at −20 °C. Over three days, six tubes were randomly selected from the freezer each day and tSGAs were extracted and quantified as outlined above by a single individual. Intraday variability was determined by computing the coefficient of variation for an analyte within a day. Interday variability was calculated by taking the coefficient of variation of all samples run over the three-day period.

### Autosampler Stability Experiments

A quality control sample containing multiple tomato species, as described, above was extracted with the addition of 100 µL of IS solution as outlined previously. Over a period of 12 hours, the quality control sample was injected and analyzed by UHPLC-MS/MS at hourly intervals. The vial cap was replaced after each injection to prevent sample evaporation between injections and the autosampler compartment was maintained at 20 °C.

## 3 Results and Discussion

### 3.1 Development of High-Throughput Extraction and UHPLC-MS/MS Quantification Methods

#### Development of high-throughput extraction method

Generally, tomato samples are pulverized in a mortar in the presence of liquid nitrogen or homogenized using a blender prior to extracting tSGAs. Tomato steroidal glycoalkaloids are considered semi-polar metabolites and are typically extracted via physical disruption in a methanolic solvent system (Ballester et al., 2016; Iijima et al., 2013, 2008; Mintz-Oron et al., 2008; Moco et al., 2006). Current methods are time consuming since each sample needs to be processed individually. Our protocol features a combined homogenization/extraction step using a Geno/Grinder system that can process up to 16 samples at once. Given a five-minute homogenization/extraction, five-minute centrifugation, and an approximately ten-minute dilution/filtration step, our extraction method can process 16 samples every 20 minutes (1.25 min/sample) making it ideal for screening large tomato populations or large sample sets of tomato products. Moreover, the tomato sample is able to stay frozen until the extraction begins which prevents potential enzymatic modification and degradation of analytes.

#### Selection of Precursor Ions

Over 100 tSGAs have been tentatively identified in tomato using high-resolution mass spectrometry and some with MS/MS fragmentation (Iijima et al., 2013, 2008; Zhu et al., 2018). However, we do not know the specific concentrations of tSGA accumulating in fruits. To study tSGAs further, quantitative analysis methods are necessary. In order to maximize the amount of tSGAs detected and separated in our method, we first compiled a target list of biologically relevant tSGAs by surveying the literature (Alseekh et al., 2015; Cichon et al., 2017; Cooperstone et al., 2017; Fujiwara et al., 2004; Hövelmann et al., 2019; Iijima et al., 2009; Zhu et al., 2018). Tomato steroidal glycoalkaloid species were prioritized based on their perceived abundance in the tomato clade, previous structural characterization, and having an established record of being impacted by or a part of biological processes such as ripening or plant defense, respectively. Using this process, 18 masses covering at least 25 different tSGA species were selected for chromatographic separation and quantification.

A 50% aqueous methanolic extract from a reference material comprised of red-ripe *Solanum lycopersicum, Solanum lycopersicum* var. *cerasiforme*, and *Solanum pimpinellifolium* fruits was used for method development on a Waters Acquity UHPLC H-Class System connected to a TQ Detector triple quadrupole mass spectrometer with electrospray ionization operated in positive ion mode. A gradient progressing from 5% to 100% acetonitrile over 15 minutes run on a Waters 2.1 × 100 mm (1.7 µm particle size) column at 0.4 mL/min was used to separate as many potential analytes as possible. Selected Ion Recordings (SIRSs) of masses of interest were utilized to identify potential tSGA species. Since only two alkaloids of interest are available commercially (alpha-tomatine and tomatidine), elution order, accurate mass, and fragmentation patterns were used to assign identity all other tSGAs. Source parameters of the MS were then adjusted to the maximize signal of both identified and tentatively identified tSGAs Those tSGAs which were readily detectable in our pooled tomato quality control samples were used in our final method. While studied more extensively than many other tSGAs, we were not able to detect and quantify beta-, gamma-, and delta-tomatine in our reference material.

#### Use of Internal Standards

We tested three, commercially available potato-derived alkaloids for their suitability as internal standards to correct for inter and intraday variability created in the MS. Alpha-solanine, alpha-chaconine, and solanidine (aglycone of alpha solanine) were selected based on their similarity in structure, ionization efficiencies and retention times to tomato-derived alkaloids. However, alpha-chaconine was excluded due to co-elution with alpha-tomatine. We determined 1.25 nmol and 22.68 pmol of alpha solanine and solanidine, respectively, should be added to each sample (41.7 femtomoles of alpha solanine and 0.756 femtomoles of solanidine on column) to achieved comparable peak areas to those observed for tSGAs and their aglycones such as tomatidine and tomatidenol (Fig. 2). Alpha-solanine and solanidine multiple reaction monitoring (MRMs) experiments were then optimized in tandem with tSGAs of interest as follows.

#### Optimization of MS parameters

Desolvation temperature, desolvation gas flow rate, and cone voltage were experimentally optimized. All other source parameters remained at their recommended default settings and are reported in the footer of (Table 1). For all experiments, vial caps were replaced after each injection to prevent any possible effects from evaporation through the pierced septa. To optimize the desolvation temperature, a 50% aqueous methanolic solution of alpha-tomatine and tomatidine was injected and desolvation temperatures ranging from 350 °C to 500 °C at 25 °C increments were tested. A 500 °C desolvation temperature resulted in the highest signal. Desolvation gas flow was tested in a similar manner starting from 600 L/hr to 1000 L/hr in 100 L/hr increments. Likewise, the 1000 L/h flow rate resulted in the most signal for both analytes. Alpha-tomatine and tomatidine were used in these experiments because of their commercial availability, their structural similarity to other tSGAs of interest, and their intended use for relative quantification of all other tSGAs and their aglycones. Finally, cone voltage was optimized by injecting a 50% aqueous methanolic extract of our tomato reference material and measuring the signal of each SIR. Cone voltages ranged from 20 to 90 V and successive injections were made in 5 V increments. Optimal cone voltages were specific to each mass and are notated in Table 1. With source parameters set to optimize the signal of all precursor ions of interest, product ion scans were then conducted to tentatively identify tSGAs and aid in the development of MRM experiments, which were ultimately used for quantification.

Since each SIR yielded multiple peaks, information from product ion scans was leveraged to determine if each peak was actually a tSGA. Product ion scan experiments were created for each mass of interest and multiple collision energies (20, 45, and 65 eV) were tested. The resulting spectra generated for each peak allowed us to eliminate peaks that were isobaric with tSGAs of interest, but had product ions inconsistent with proposed structures. Masses such as 254.9 and 272.9 *m*/*z* were particularly useful in identifying alkaloids as they are likely derived from the fragmentation of the steroidal backbone characteristic of all tSGAs (Supplementary Information) and have been previously reported in the literature (Caprioli et al., 2014; Cichon et al., 2017; Iijima et al., 2013; Sonawane et al., 2018; Zhu et al., 2018). Additionally, tSGAs with the prefix “dehydro” exhibit a desaturation on the B ring of the steroidal backbone between carbons 5 and 6 (Iijima et al., 2013; Itkin et al., 2011; Ono et al., 1997; Sonawane et al., 2018). We observed that common fragments derived from the steroidal backbone of these alkaloids, such as 252.9 and 270.8, were accordingly 2 *m*/*z* less than their saturated counterparts. The 272.9 fragment corresponds to the A-D rings of the steroidal backbone and its corresponding water loss product (Sonawane et al., 2018). Elution order of analytes was used to help tentatively identify tSGAs detected in our reference sample based on previous reports (Alseekh et al., 2015; Zhu et al., 2018). Multiple collision energies allowed us to select product ions that were abundant and consistently produced under different conditions. These product ions then became candidate ions for MRM development.

MRM experiments allowed us to confidently detect and quantify tSGAs of interest and increase sensitivity by minimizing interference of co-eluting compounds. We created MRM experiments for each mass using optimized source conditions and four product ions with the highest signal/noise ratio. Initially, our 50% aqueous methanolic reference sample extract was injected and each transition was tested at 5 eV. The experiments were rerun at increasing collision energies at 15 eV increments up to 95 eV. Afterwards, a 20 eV window broken into 5 eV increments was determined for each transition and the experiments were re-run. Optimized MRMs are displayed in Table 1. To maximize duty cycle, two transitions with the best signal to noise ratio were retained. The gradient was then optimized to chromatograph each analyte. All tSGAs were quantified using a standard curve generated with alpha-tomatine while aglycone species used tomatidine. Due to the structural similarity among tSGA species quantified in our method, we hypothesize that ionization efficiencies will be similar amongst our analytes. Lastly, MRMs were developed for the potato derived alkaloids alpha-solanine and solanidine used as IS. These IS allowed us to correct for instrument derived variability that normally occurs with mass spectrometers.

#### Development of Chromatographic Gradient

Method development related to the MS was initially carried out using a simple 13-minute gradient outlined above. While this run time is shorter than many of the previously published studies characterizing tSGAs using high-resolution MS (Iijima et al., 2013, 2008; Zhu et al., 2018) we aimed to create a more efficient method that would be able to accommodate large sample sets. Of the two columns tested (Waters C18 Acquity bridged ethylene hybrid (BEH) 2.1 × 100 mm, 1.7 µm and Waters C18 Acquity high strength silica (HSS) 2.1 × 100 mm, 1.8 µm), the BEH column was able to better resolve analytes of interest with a particular benefit observed in the nonpolar aglycone steroidal alkaloids. We adjusted our gradient conditions in such a way that all separation of analytes occurred within a six-minute window with an additional five minutes devoted to cleaning and requilibrating the column to reduce carryover (Fig. 2). Additionally, the needle wash was set to rinse the needle and injection port for ten seconds before and after an injection with 1:1 methanol:isopropanol to further reduce carryover. We observed multiple peaks for many of our masses indicating the presence of multiple isobaric tSGAs (likely including structural isomers) (Fig. 2). In the case of esculeoside B, multiple diastereomers have been previously reported in tomato products which explains our observation of multiple peaks for this analyte (Hövelmann et al., 2019; Manabe et al., 2013; Nohara et al., 2015). Validation experiments, including confirmation of peak identities using high-resolution mass spectrometry, were next carried out using the finalized chromatographic gradient.

### 3.2 Validation of Extraction and UHPLC-MS/MS Methods

#### Confirmation of Analytes using High-Resolution Mass Spectrometry

Accurate mass spectrometry was used to confirm the identities of analytes quantified by our UHPLC-MS/MS method. We transferred our method to an Agilent 1290 Infinity II connected to an Agilent 6545 QTOF and profiled tSGAs both in high resolution full scan mode (50-1700 *m*/*z*) and through targeted fragmentation experiments. Both types of experiments were consistent with our identities of all tSGAs and aglycones in our UHPLC-MS/MS method (Table 2). Retention times differed slightly between the UHPLC-MS/MS method and the UHPLC-QTOF-MS experiments due to differences in dead volume between the two instruments. However, relative elution order remained the same.

**Table 2.** UHPLC-QTOF-MS Confirmation of tSGA Identities

| Tentative Identification | Molecular Formula | Retention Time (min) | Monoisotopic Mass | Observed Mass [M+H] | Mass Error ($\Delta$ ppm) | Common MS/MS Fragments<sup>a</sup> |
| --- | --- | --- | --- | --- | --- | --- |
| Esculeoside B | C<sub>56</sub>H<sub>93</sub>NO<sub>28</sub> | 2.24 | 1227.5884 | 1228.5989 | 2.20 | 1048.5380, 273.2120, 255.2016, 163.0509, 145.0404, 85.0205 |
|  |  | 2.34 |  | 1228.5967 | 0.41 |  |
|  |  | 2.45 |  | 1228.5966 | 0.33 |  |
| Hydroxytomatine | C<sub>50</sub>H<sub>83</sub>NO<sub>22</sub> | 2.84 | 1049.5407 | 1050.5500 | 1.43 | 1032.5385, 273.2213, 255.2203, 161.1318, 145.0489, 85.0279 |
|  |  | 3.22 |  | 1050.5513 | 2.67 |  |
|  |  | 3.29 |  | 1050.5506 | 2.00 |  |
|  |  | 3.50 |  | 1050.5501 | 1.52 |  |
| Dehydrolycoperoside F, G, or Dehydroesculeoside A | C<sub>58</sub>H<sub>93</sub>NO<sub>29</sub> | 2.41 | 1267.5828 | 1268.5930 | 1.89 | 1208.5714, 1046.5175, 271.2054, 253.1951, 163.0600, 85.0284 |
| Lycoperoside F, G, or Esculeoside A | C<sub>58</sub>H<sub>95</sub>NO<sub>29</sub> | 2.44 | 1269.5985 | 1270.6076 | 1.02 | 1210.5900, 1048.5324, 273.2213, 255.2108, 163.0600, 85.0285 |
|  |  | 3.06 |  | 1270.6095 | 2.52 |  |
| Acetoxytomatine (I) | C<sub>52</sub>H<sub>85</sub>NO<sub>23</sub> | 4.22 | 1091.5507 | 1092.5614 | 2.65 | 1032.5386, 273.2216, 255.2112, 161.1326, 145.0497, 85.0287 |
| Dehydrotomatine<sup>b</sup> | C<sub>50</sub>H<sub>81</sub>NO<sub>21</sub> | 4.99 | 1031.5301 | 1032.5388 | 0.87 | 1014.5274, 271.2054, 253.1951, 145.0495, 85.0284, 57.0337 |
|  |  | 5.20 |  | 1032.5373 | 0.58 |  |
| Acetoxytomatine (II) | C<sub>52</sub>H<sub>85</sub>NO<sub>23</sub> | 5.32 | 1091.5507 | 1092.5619 | 3.11 | 1032.5404, 273.2216, 255.2114, 161.1328, 145.0499, 85.0288 |
|  |  | 5.39 |  | 1092.5608 | 2.11 |  |
| Alpha-tomatine<sup>b</sup> | C<sub>50</sub>H<sub>83</sub>NO<sub>21</sub> | 5.35 | 1033.5457 | 1034.5557 | 2.13 | 1016.5449, 416.3523, 273.2217, 255.2112, 145.0498, 85.0287 |
| Tomatidine<sup>b</sup> | C<sub>27</sub>H<sub>45</sub>NO<sub>2</sub> | 6.95 | 415.3450 | 416.3531 | 0.72 | 398.3414, 273.2208, 255.2101, 161.1318, 126.1271, 81.0693 |
| Tomatidenol<sup>b</sup> | C<sub>27</sub>H<sub>43</sub>NO<sub>2</sub> | 6.98 | 413.3294 | 414.3371 | 0.24 | 396.3260, 271.2053, 253.1949,<br>161.1322, 126.1275, 81.0695 |

<sup>a</sup>MS/MS product ions generated at 70 eV and 45 eV for glycosylated and aglycone species, respectively. Other source parameters
were previously enumerated.
<sup>b</sup>Identification confirmed by authentic standard

Targeted MS/MS experiments using the UHPLC-QTOF-MS allowed us to determine common spectral characteristics for each tSGA (Fig. 1 and Supplementary Information). Using commercially available alpha-tomatine and tomatidine and exploiting the presence of dehydrotomatine and tomatidenol (dehydrotomatine) as impurities within these standards, we were able to collect MS/MS fragmentation data on these four analytes. We found that all tSGAs and aglycones fragmented in predictable ways that allow for identification. Common masses produced by each tSGA in our method can be found in Table 2. These data allow us to tentatively identify all analytes in our UHPLC-MS/MS with a high degree of confidence.

#### LOD and LOQ

Previous chromatography-based methods to quantify both potato and tSGAs relied on photodiode array detectors and set 208 nm (Del Giudice et al., 2015; Kozukue et al., 2004; Kozukue and Friedman, 2003; Tajner-Czopek et al., 2014). Given that the molar extinction coefficient for alpha-tomatine is only 5000 M^-1^c^-1^, (Keukens et al., 1994), photodiode array detectors are not sensitive enough for detecting low quantities of these compounds, nor distinguishing between different alkaloids. Moreover, photodiode array detectors are often set to 200 nm to quantify tSGAs which is a non-specific wavelength where many compounds (including mobile phases) can absorb light (Friedman and Levin, 1998, 1992; Keukens et al., 1994). Mass spectrometers offer substantial gains in sensitivity through the use of MRM experiments and the ability to differentiate numerous analytes in a single run. Our UHPLC-MS/MS method for quantifying tSGAs was able to detect and quantify alpha-tomatine and tomatidine in the low femtomole-on-column range (Table 3). Given our extraction method, tSGAs could be present in picomolar concentrations in tomato and still be quantified.

**Table 3.** Extraction efficiency of commercially available tSGAs and potato-derived internal standards.

| Analyte | Sample Size | Extraction Efficiency (%) | LOD (femtomoles injected) | LOQ (femtomoles injected) |
| --- | --- | --- | --- | --- |
| Alpha-tomatine | n=6 | 100.8 ± 13.1 | 1.0988 | 0.3296 |
| Alpha-solanine<sup>a</sup> | n=6 | 94.3 ± 3.4 | N/A<sup>b</sup> | N/A |
| Tomatidine | n=6 | 93.0 ± 6.8 | 0.3354 | 0.1006 |
| Solanidine<sup>a</sup> | n=6 | 99.7 ± 7.1 | N/A | N/A |

<sup>a</sup>Analyte used as an internal standard with no calibration curve
<sup>b</sup>Not applicable due to its use as an internal standard

#### Spike Recovery

Spike addition experiments were conducted to assess the performance of our high-throughput extraction method. Both tomato and potato derived external alkaloid standards were used to determine if our chosen internal standards would behave similarly to analytes native to tomato. Tomato alkaloids alpha-tomatine (100.8% ± 13.1) and tomatidine (93% ± 6.8) as well as the potato-derived internal standards alpha solanine (94.3% ± 3.4) and solanidine (99.7% ± 7.1) were efficiently extracted using our method (Table 3). These data indicate that our method is able to effectively extract aglycone and glycosylated steroidal alkaloid species from tomato and our internal standards extract similarly to native analytes.

#### Intra/Interday Variability

Experiments to determine intra/interday variability were conducted to determine analytical variability in our extraction and analysis methods. A single operator extracted six tomato samples and analyzed them by UHPLC-MS/MS. This experiment was repeated twice more by the same operator. Our data indicate that our methods are reliable with most analytes having coefficient of variations for both intra and interday variability below 5% (Table 4). As expected, interday variability was higher than intraday variability for all analytes reflecting day-to-day variability in the MS.

**Table 4.**
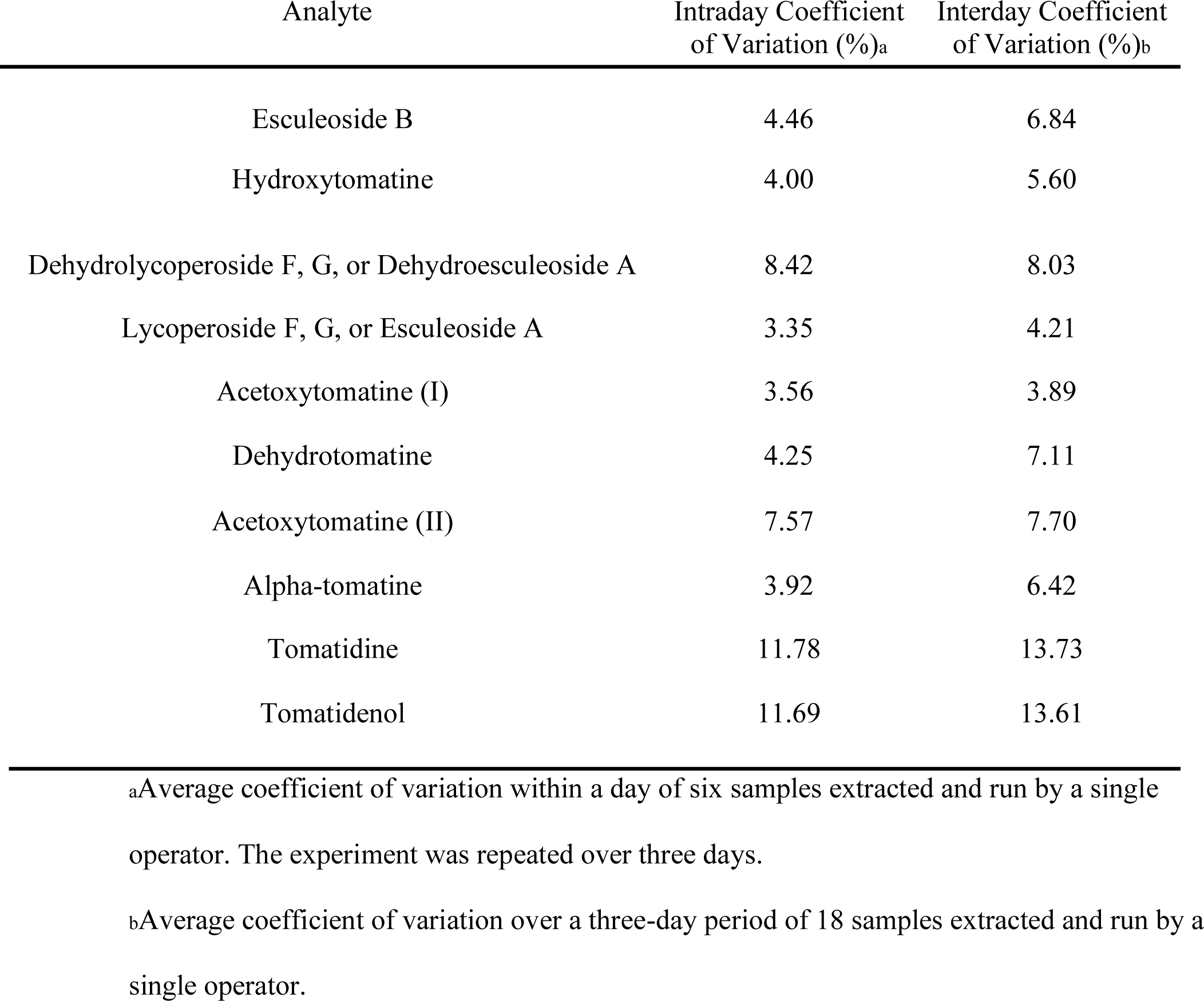
Intraday and interday coefficient of variation values for analytes quantified by our UHPLC-MS/MS method

| Analyte | Intraday Coefficient<br>of Variation (%)<sup>a</sup> | Interday Coefficient<br>of Variation (%)<sup>b</sup> |
| --- | --- | --- |
| Esculeoside B | 4.46 | 6.84 |
| Hydroxytomatine | 4.00 | 5.60 |
| Dehydrolycoperoside F, G, or Dehydroesculeoside A | 8.42 | 8.03 |
| Lycoperoside F, G, or Esculeoside A | 3.35 | 4.21 |
| Acetoxytomatine (I) | 3.56 | 3.89 |
| Dehydrotomatine | 4.25 | 7.11 |
| Acetoxytomatine (II) | 7.57 | 7.70 |
| Alpha-tomatine | 3.92 | 6.42 |
| Tomatidine | 11.78 | 13.73 |
| Tomatidenol | 11.69 | 13.61 |

<sup>a</sup>Average coefficient of variation within a day of six samples extracted and run by a single operator. The experiment was repeated over three days.
<sup>b</sup>Average coefficient of variation over a three-day period of 18 samples extracted and run by a single operator.

#### 12-hour Stability Experiment

Tomato phytochemicals typically analyzed, such as carotenoids, are subject to oxidation and need to be run in small batches to minimize experimental error due to degradation (Kopec et al., 2012). However, relatively little is known about the stability of tSGAs compared to the above phytochemical classes. We hypothesized that due to the known heat stability of chemically analogous potato steroidal glycoalkaloids, extracted tSGAs would be stable over time. A 12-hour stability study demonstrated that both alpha-tomatine and tomatidine did not degrade over time in an autosampler maintained at 20 °C. While there is currently no published literature investigating the stability of tSGAs, some data exists in chemically analogous potato glycoalkaloids. Often, potato glycoalkaloids are often extracted at 100 °C temperatures to disrupt cell walls and otherwise weaken the sample matrix (Rodriguez-Saona et al., 1999) and processing studies have shown that these compounds are stable up to 180 °C (Chungcharoen, 1988). Therefore, tSGAs may also have similar heat tolerance attributes and we speculate that these analytes may remain unchanged in autosamplers well beyond the 12-hour time period we tested.

### 3.3 Application of Extraction and UHPLC-MS/MS Method

#### Grocery Store Survey

To test our extraction and quantification method, we surveyed several commonly consumed tomato-based products available at grocery stores. The purpose was twofold: to test applicability of or method, and to report comprehensive and quantitative values of tSGAs in commonly consumed tomato products. These products included an assortment of fresh tomatoes, ketchup, pasta sauce, pizza sauce, tomato soup, tomato paste, tomato juice, and whole peeled tomatoes (Table 5). Values are reported per serving to normalize between tomato products subjected to varying degrees of concentration. While there are some reports of tSGA concentrations in fresh tomatoes using modern methods (Baldina et al., 2016), concentrations in tomato-based products are not well reported in the literature. We found that tSGAs varied depending on type of product. High standard deviations likely reflect differences in geographic origin, harvest time, and processing conditions. Of note, many of our tSGAs varied by up to three orders of magnitude among different analytes and tomato products. This finding indicates a broad range of tSGA concentrations in tomato-based products.

**Table 5.**
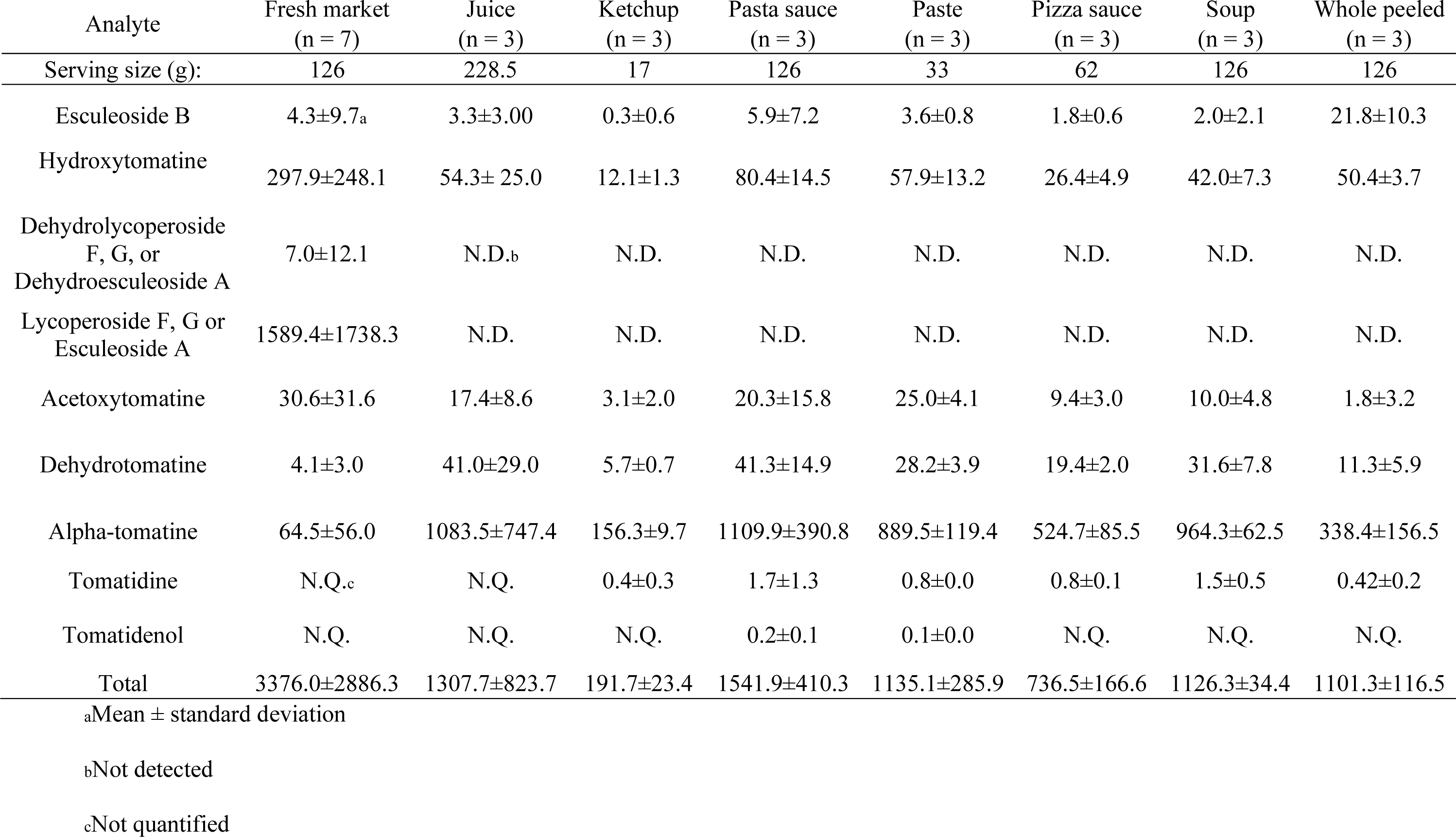
Survey of tSGAs in common tomato-based products reported in µg per serving size

| Analyte | Fresh market<br>(n = 7) | Juice<br>(n = 3) | Ketchup<br>(n = 3) | Pasta sauce<br>(n = 3) | Paste<br>(n = 3) | Pizza sauce<br>(n = 3) | Soup<br>(n = 3) | Whole peeled<br>(n = 3) |
| --- | --- | --- | --- | --- | --- | --- | --- | --- |
| Serving size (g): | 126 | 228.5 | 17 | 126 | 33 | 62 | 126 | 126 |
| Esculeoside B | 4.3±9.7<sub>a</sub> | 3.3±3.00 | 0.3±0.6 | 5.9±7.2 | 3.6±0.8 | 1.8±0.6 | 2.0±2.1 | 21.8±10.3 |
| Hydroxytomatine | 297.9±248.1 | 54.3± 25.0 | 12.1±1.3 | 80.4±14.5 | 57.9±13.2 | 26.4±4.9 | 42.0±7.3 | 50.4±3.7 |
| Dehydrolycoperoside<br>F, G, or<br>Dehydroesculeoside A | 7.0±12.1 | N.D.<sub>b</sub> | N.D. | N.D. | N.D. | N.D. | N.D. | N.D. |
| Lycoperoside F, G or<br>Esculeoside A | 1589.4±1738.3 | N.D. | N.D. | N.D. | N.D. | N.D. | N.D. | N.D. |
| Acetoxytomatine | 30.6±31.6 | 17.4±8.6 | 3.1±2.0 | 20.3±15.8 | 25.0±4.1 | 9.4±3.0 | 10.0±4.8 | 1.8±3.2 |
| Dehydrotomatine | 4.1±3.0 | 41.0±29.0 | 5.7±0.7 | 41.3±14.9 | 28.2±3.9 | 19.4±2.0 | 31.6±7.8 | 11.3±5.9 |
| Alpha-tomatine | 64.5±56.0 | 1083.5±747.4 | 156.3±9.7 | 1109.9±390.8 | 889.5±119.4 | 524.7±85.5 | 964.3±62.5 | 338.4±156.5 |
| Tomatidine | N.Q.<sub>c</sub> | N.Q. | 0.4±0.3 | 1.7±1.3 | 0.8±0.0 | 0.8±0.1 | 1.5±0.5 | 0.42±0.2 |
| Tomatidenol | N.Q. | N.Q. | N.Q. | 0.2±0.1 | 0.1±0.0 | N.Q. | N.Q. | N.Q. |
| Total | 3376.0±2886.3 | 1307.7±823.7 | 191.7±23.4 | 1541.9±410.3 | 1135.1±285.9 | 736.5±166.6 | 1126.3±34.4 | 1101.3±116.5 |

<sub>a</sub>Mean ± standard deviation
<sub>b</sub>Not detected
<sub>c</sub>Not quantified

Alpha-tomatine, the first tSGA in the biosynthesis pathway, was found to be in the highest concentration in processed tomato products such as paste, pasta sauce, and soup (Table 5). The discrepancy between fresh and whole peeled tomatoes is hypothesized to be due to genetic and environmental conditions that influenced the chemical profile of the tomatoes prior to processing. Analyte groups like dehydrolycoperoside F, G or A, lycoperosides F, G, or esculeoside A and acetoxytomatine (commonly referred to as lycoperosides A, B, or C) were not detectable in most tomato products except for some fresh varieties and ketchup. Interestingly, lycoperosides F, G, or esculeoside A are typically the most abundant tSGA in fresh tomatoes. This observation raises questions about the effects of processing on tSGAs where few studies have been conducted to date (Tomas et al., 2017). While the chemically analogous potato glycoalkaloids are considered to be heat stable, high temperatures, pressures, and any combination thereof might be detrimental to some tSGAs or cause shifts in chemical profiles.

Concentrations of tSGAs in tomato products were normalized for serving size to contextualize how much might be ingested in a given meal. Other tomato phytochemicals, such as lycopene, tend to be found in concentrations ranging from 0.09 to 9.93 mg/100g FW in fresh tomatoes (Dzakovich et al., 2019). Compared to major carotenoids found in tomato, tSGA concentrations were comparable (0.7 to 3.4 mg/serving) (Cooperstone, 2020) This finding contradicts a long-standing misconception that tSGAs are degraded during ripening (Friedman, 2002). Rather, tSGAs such as alpha-tomatine are biochemically transformed during ripening into glycosylated and acetylated forms. Overall, our methods were able to efficiently extract and analyze many types of tSGAs and generate the first quantitative concentration reports of these analytes in commonly consumed tomato products. Moreover, we found that tSGAs can be found in similar concentrations to other major phytochemicals in tomatoes such as carotenoids.

We have developed and described the first comprehensive extraction and analysis method for tSGAs. Our extraction method was able to quickly and efficiently extract tSGAs and allowed for high-throughput workflows (16 samples per ∼20 min) to be utilized. Our UHPLC-MS/MS method was able to separate and quantify 16 tSGAs representing 9 different tSGA masses, as well as two internal standards, in 13 minutes. Limits of quantification for commercially available tSGAs were 1.09 and 0.34 femtomoles on column for alpha-tomatine and tomatidine, respectively. This corresponds to 0.8 and 0.25 µg/100g of alpha-tomatine and tomatidine in tomato, respectively, given our extraction procedures. Relative quantification for tSGAs and aglycones that did not have commercially available standards was performed using alpha-tomatine and tomatidine, respectively. Our methods were able to successfully profile tSGAs in a comprehensive array of commonly available tomato-based products. These values are among the first to be reported in the literature and can serve as benchmarks for future studies investigating tSGAs in a variety of contexts. Our extraction and UHPLC-MS/MS method will allow researchers to rapidly and accurately generate data about tSGAs and overcomes a major limitation hampering this field and allow for the field to advance.

## Acknowledgements

We thank David Francis, Jiheun Cho, Troy Aldrich (The Ohio State University, Ohio Agriculture Research and Development Center), and the North Central Agricultural Research Station crews for assistance with selecting, planting, and harvesting tomatoes used in this study.

## Data Availability Statement

The raw data supporting the conclusions of this manuscript will be made available by the authors, without undue reservation, to any qualified researchers.

