## Supplementary figures 1-9 for "A High-Throughput Extraction and Analysis Method for Steroidal Glycoalkaloids in Tomato"

Supplementary Material

**Supplementary Figure 1.** Fragmentation patterns for esculeoside B from a tomato quality control sample generated with UHPLC-QTOF-MS. The instrument was run in targeted MS/MS mode with a collision energy of 70 eV.

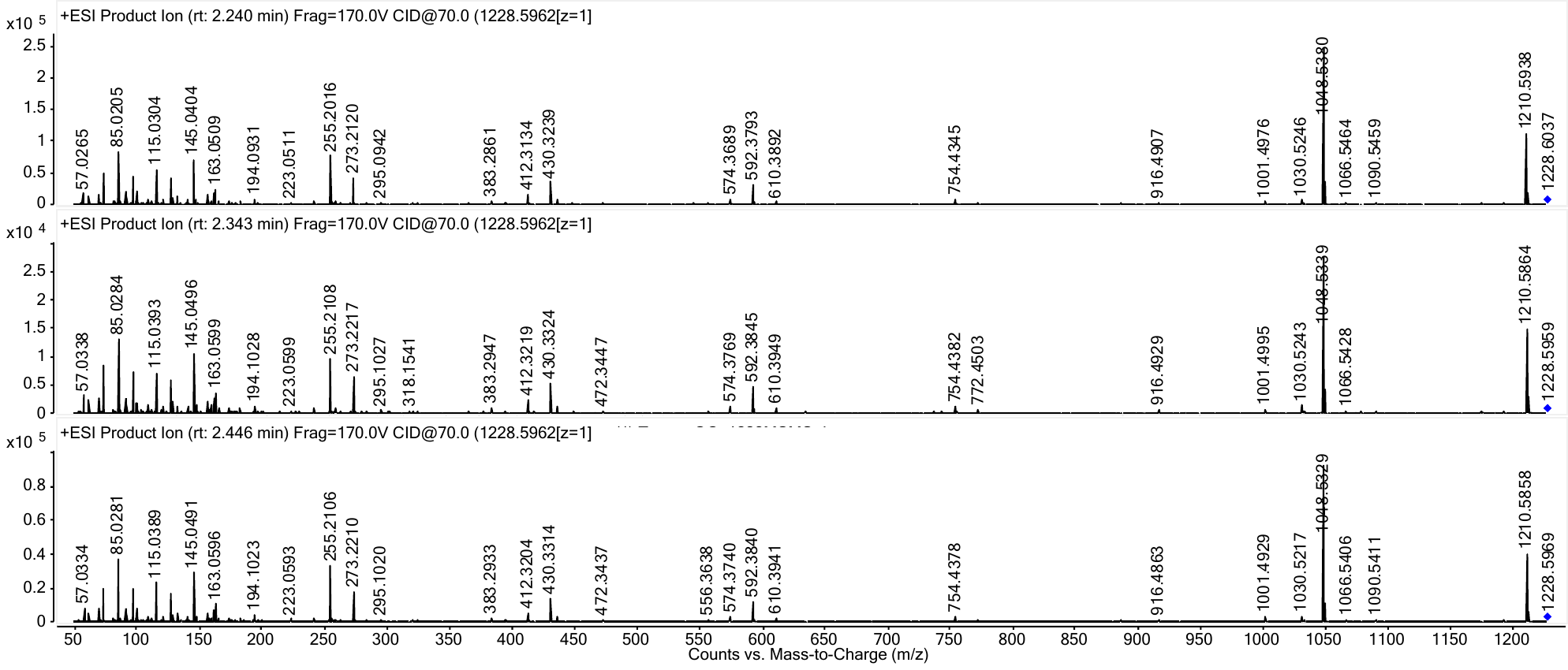

**Supplementary Figure 2.** Fragmentation patterns for hydroxytomatine from a tomato quality control sample generated with UHPLC-QTOF-MS. The instrument was run in targeted MS/MS mode with a collision energy of 70 eV.

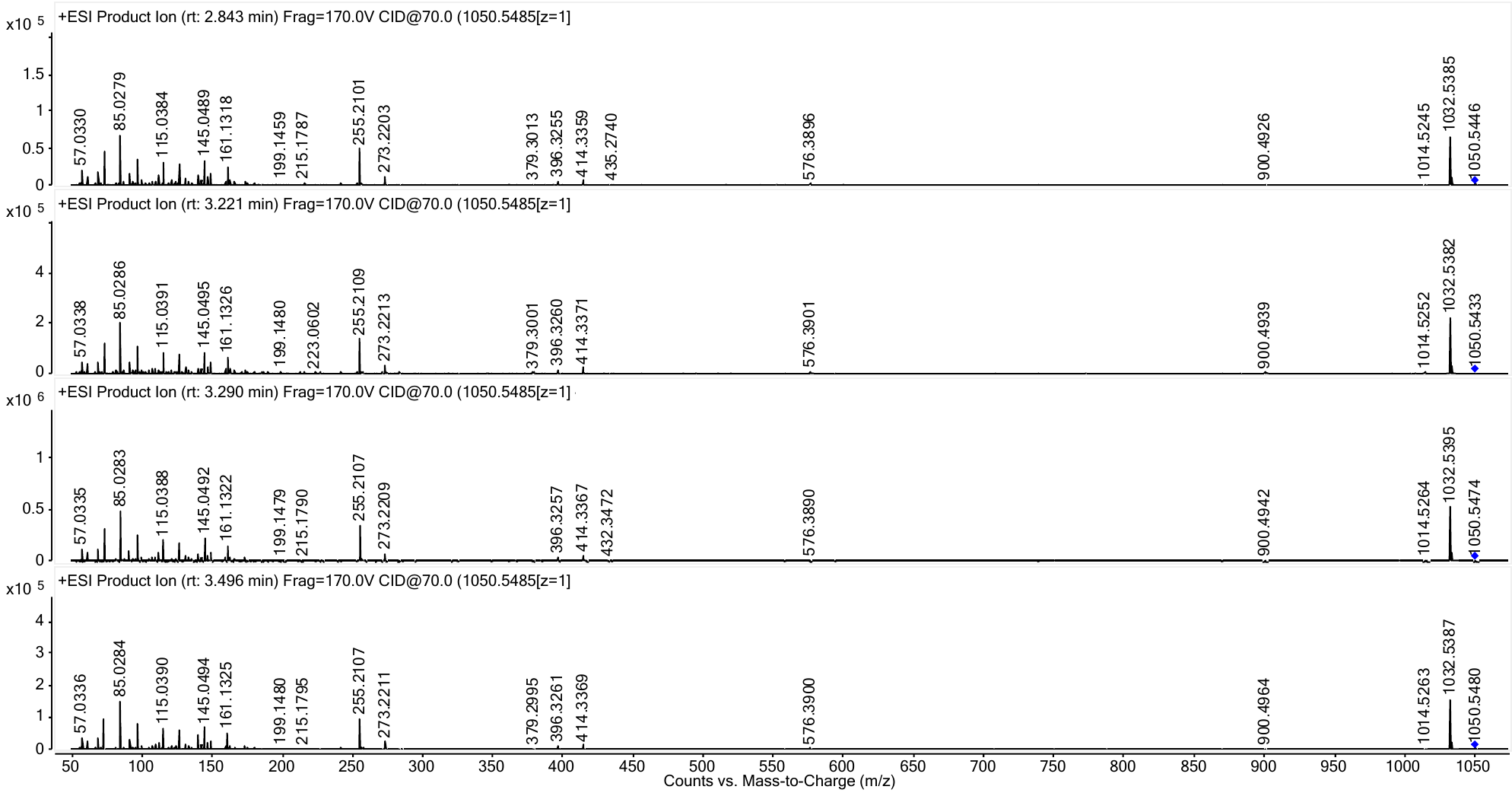

**Supplementary Figure 3.** Fragmentation patterns for dehydrolycoperoside F, G, or dehydroesculeoside A from a tomato quality control sample generated with UHPLC-QTOF-MS. The instrument was run in targeted MS/MS mode with a collision energy of 70 eV.

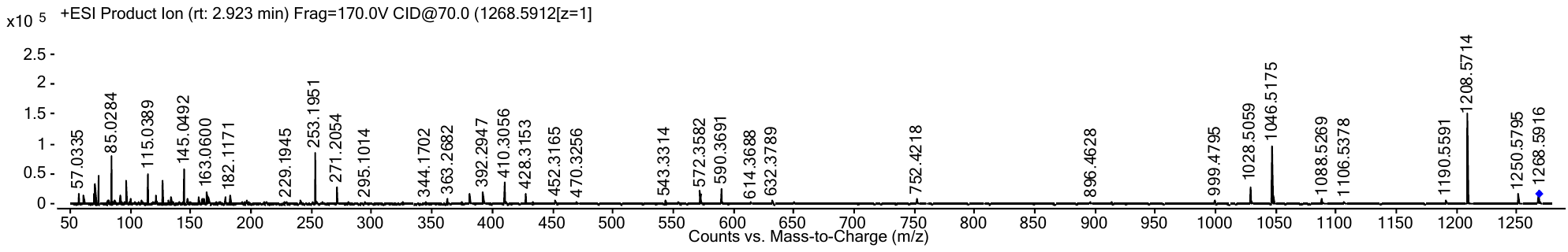

**Supplementary Figure 4.** Fragmentation patterns for lycoperoside F, G, or esculeoside A from a tomato quality control sample generated with UHPLC-QTOF-MS. The instrument was run in targeted MS/MS mode with a collision energy of 70 eV.

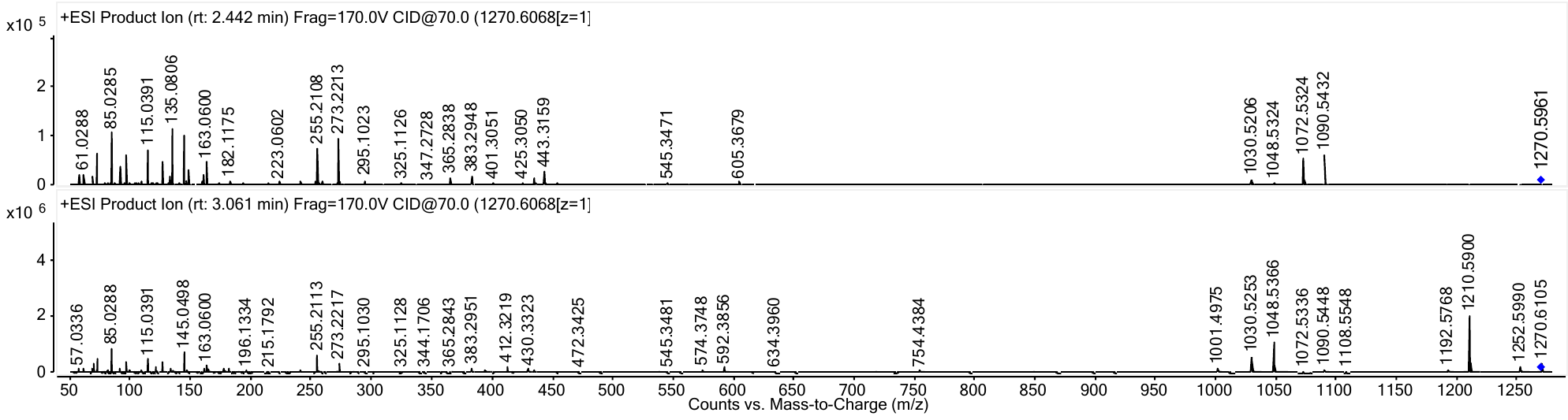

**Supplementary Figure 5.** Fragmentation patterns for acetoxytomatine (also referred to as lycoperosides A, B, and C) from a tomato quality control sample generated with UHPLC-QTOF-MS. The instrument was run in targeted MS/MS mode with a collision energy of 70 eV.

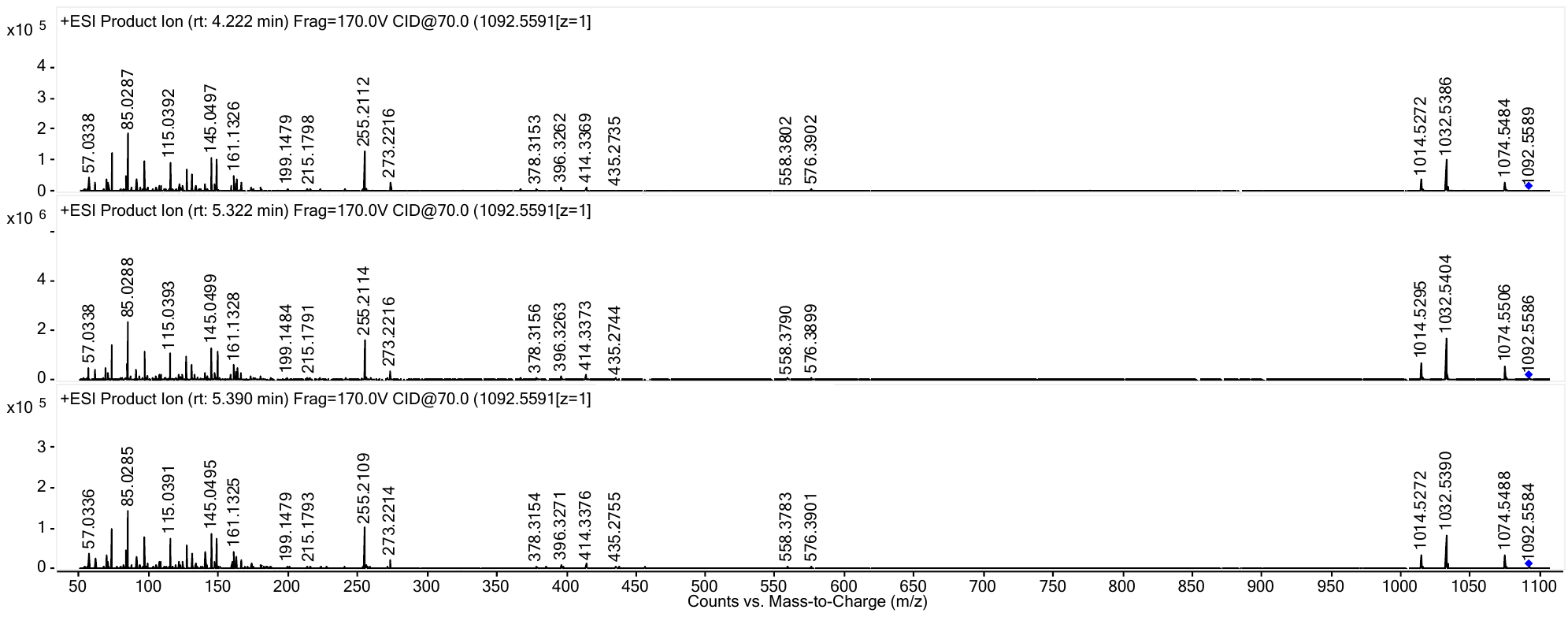

**Supplementary Figure 6.** Fragmentation patterns for alpha-tomatine from a tomato quality control sample (A) and an authentic standard (B) generated with UHPLC-QTOF-MS. The instrument was run in targeted MS/MS mode with a collision energy of 70 eV.

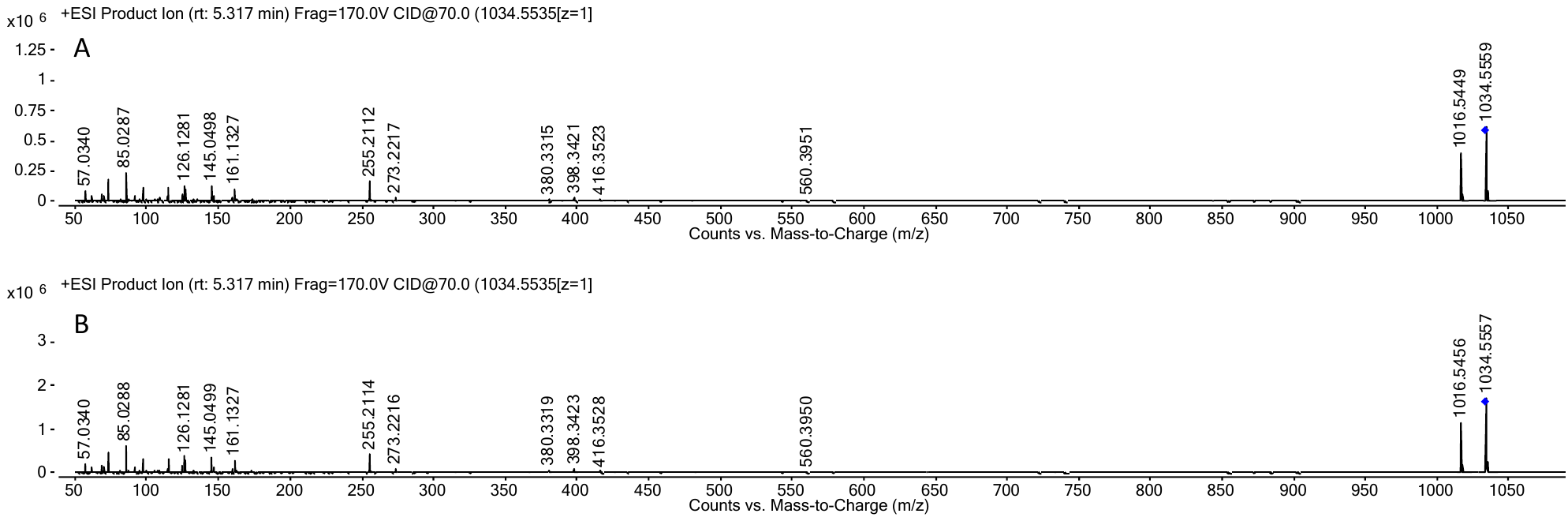

**Supplementary Figure 7.** Fragmentation patterns for dehydrotomatine from a tomato quality control sample (A) and an authentic standard (B) generated with UHPLC-QTOF-MS. The instrument was run in targeted MS/MS mode with a collision energy of 70 eV.

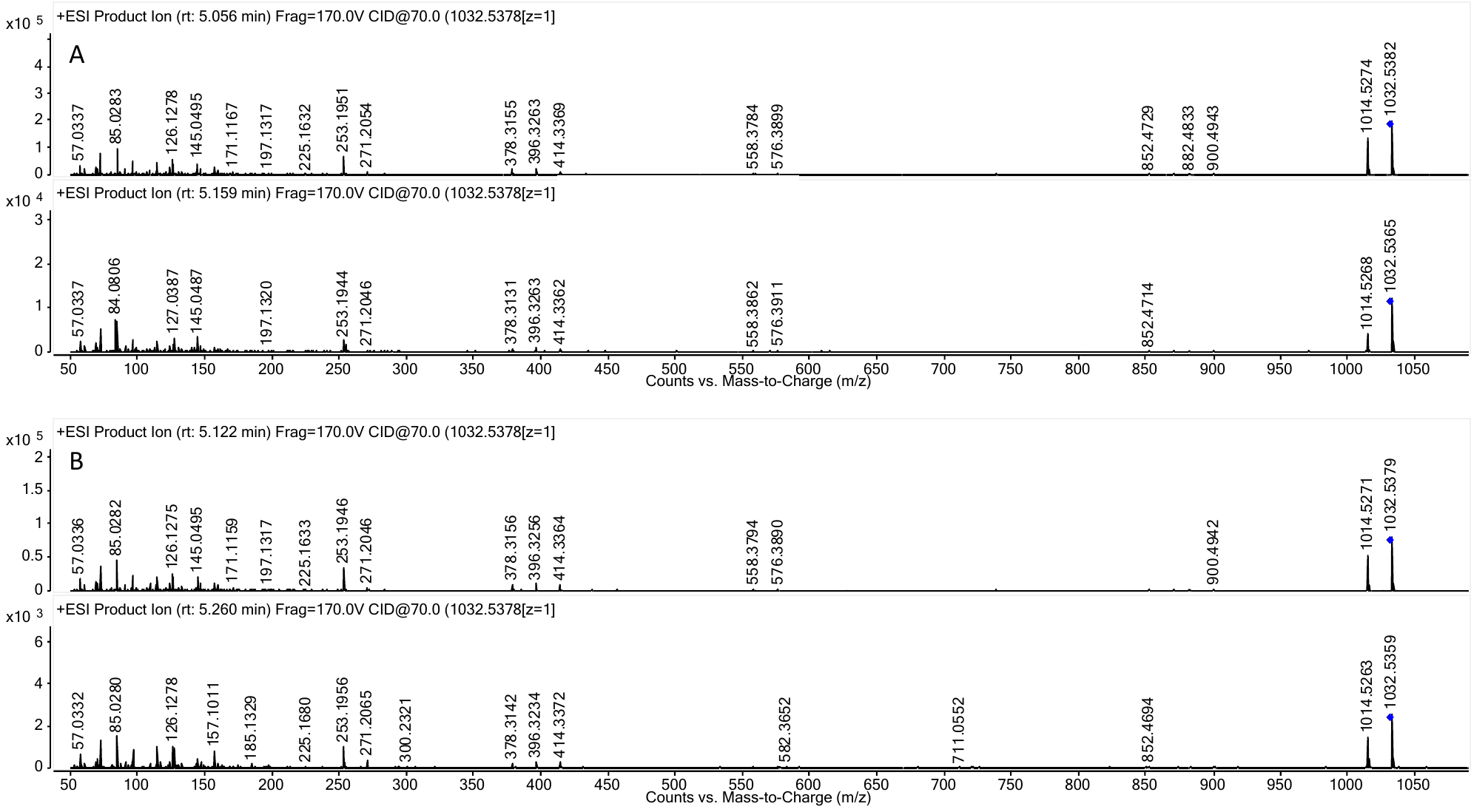

**Supplementary Figure 8.** Fragmentation patterns for tomatidine from a tomato quality control sample (A) and an authentic standard (B) generated with UHPLC-QTOF-MS. The instrument was run in targeted MS/MS mode with a collision energy of 45 eV.

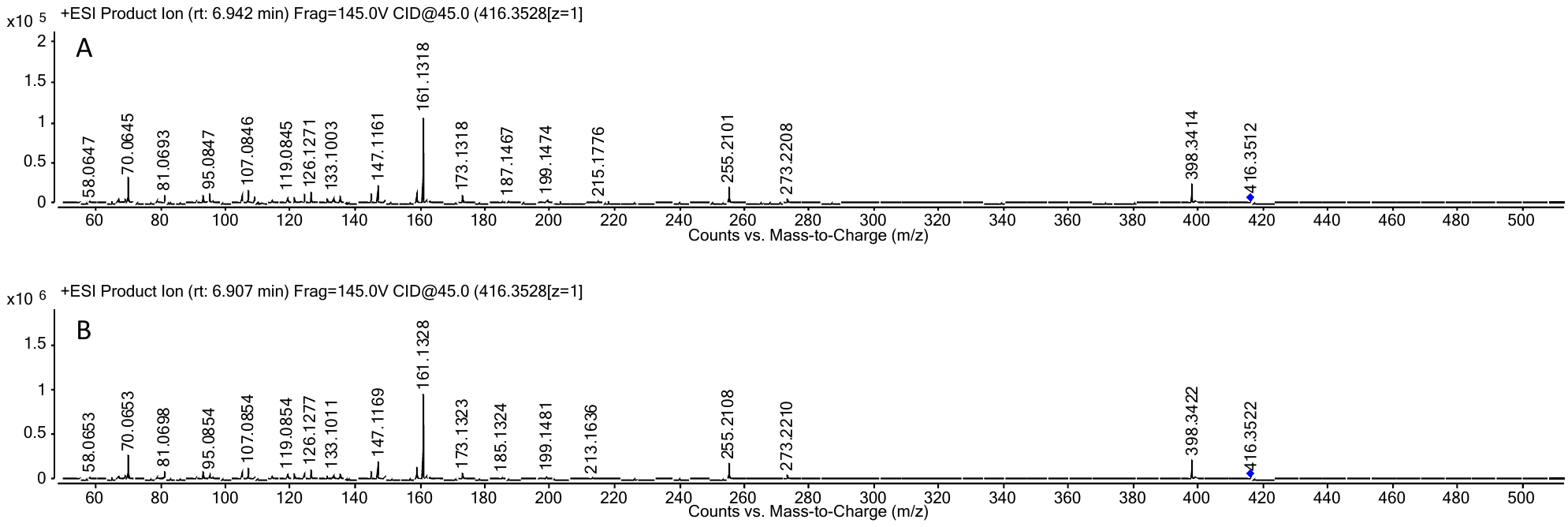

**Supplementary Figure 9.** Fragmentation patterns for tomatidenol (dehydrotomatidine) from a tomato quality control sample (A) and an authentic standard (B) generated with UHPLC-QTOF-MS. The instrument was run in targeted MS/MS mode with a collision energy of 45 eV.

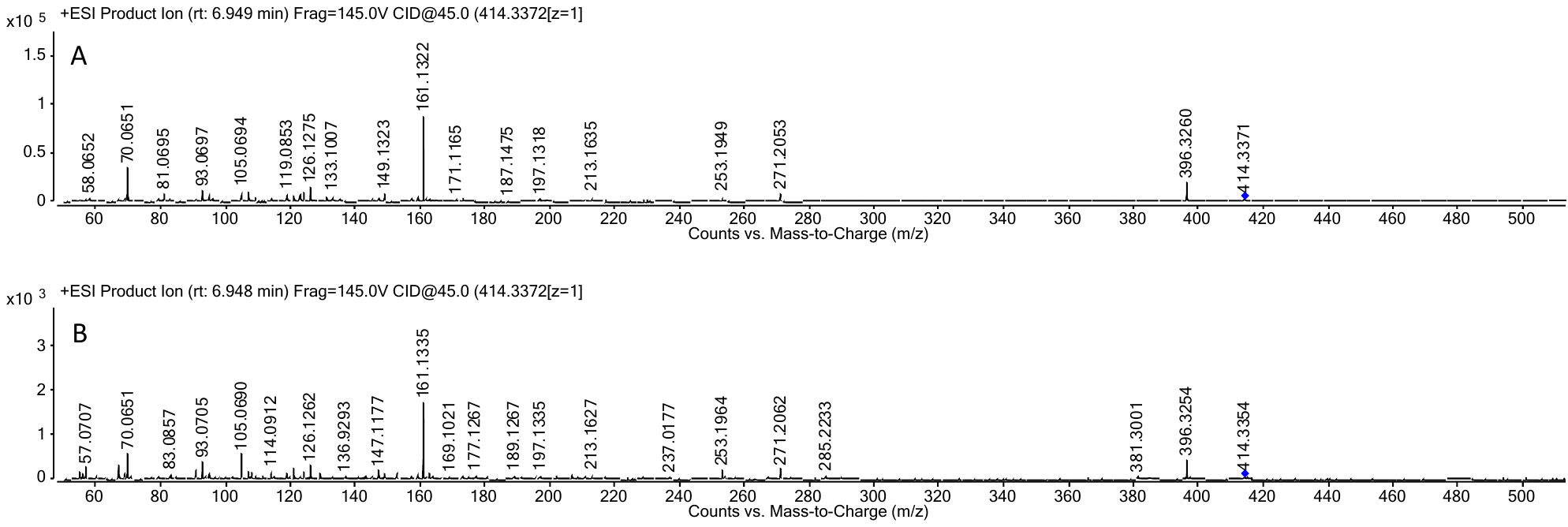
